# Development of the alpha rhythm is linked to visual white matter pathways and visual detection performance

**DOI:** 10.1101/2022.09.03.506461

**Authors:** Sendy Caffarra, Klint Kanopka, John Kruper, Adam Richie-Halford, Ethan Roy, Ariel Rokem, Jason D. Yeatman

## Abstract

Alpha is the strongest electrophysiological rhythm in awake humans at rest. Despite its predominance in the EEG signal, strong variations can be observed in alpha properties during development, with an increase of alpha frequency over childhood and adulthood. Here we tested the hypothesis that these changes of alpha rhythm are related to the maturation of visual white matter pathways. We capitalized on a large dMRI-EEG dataset (dMRI n=2,747, EEG n=2,561) of children and adolescents (age range: 5-21 years old) and showed that maturation of the optic radiation specifically accounts for developmental changes of alpha frequency. Behavioral analyses also confirmed that variations of alpha frequency are related to maturational changes in visual perception. The present findings demonstrate the close link between developmental variations in white matter tissue properties, electrophysiological responses, and behavior.

## Introduction

The alpha rhythm (8-12 Hz) is one of the most prominent and consistent electrophysiological brain signatures in both human and animal brains^1–3^. In the 1920s, this dominant brain rhythm was first reported in humans at rest by Hans Berger^1^. Despite its widespread occurrence and long history, the neuroanatomical structures that influence the alpha rhythm and its development are still under discussion. This work capitalizes on structural (diffusion magnetic resonance imaging; dMRI) and functional (electroencephalography; EEG) brain measures from a large sample (dMRI n=2,747, EEG n=2,561) spanning 5 to 21 years of age to clarify the neurobiological underpinnings of human spontaneous alpha across development.

One of the major neural sources of the alpha rhythm is the thalamus (pulvinar and lateral geniculate nucleus, LGN^4–6^), whose crucial role has been shown in in-vitro slice preparations^7^ and further confirmed by human studies where fluctuations in thalamic activity (due to tasks or lesions) lead to changes in occipital alpha^8–10^. Besides the thalamus, additional alpha generators have been localized in the visual cortex of both humans and animals^2,11,12^. Crucially, these thalamic and cortical generators can synchronize and show a high degree of alpha coherence^13,14^. This supports the idea that modulations of alpha depend not only on a single brain area’s activity but also on cortico-thalamic connections^15^ and their white matter properties^16^. Specifically, mathematical models of spontaneous brain rhythms have proposed that electrophysiological oscillations can be described as a function of the structural properties of white matter fibers (e.g., fiber length, localization, distribution, density;^17–19^). Despite the high precision of these models, there is still no definitive evidence supporting the theorized link between white matter and alpha. The optic radiation has been the most studied cortico-thalamic pathway since it connects two major alpha generators: LGN and the primary visual cortex^20^. However, the experimental findings are mixed, with research studies reporting positive effects^21,22^, null effects^23^ and studies highlighting the role of other cortical connections (corona radiata, corpus callosum^24^). Part of the source of this incongruence likely stems from small sample sizes: most previous studies linking alpha to white matter properties included 20-30^20,21,22^ participants and the largest study to date included 89 participants^24^.

The present work leverages a large EEG-dMRI dataset (dMRI n=2,747, EEG n=2,561) including children and adults ranging from 5 to 21 years of age and reveals a relationship between white matter fiber properties and individual differences in spontaneous alpha activity. This structural-functional link is specific to the optic radiation, and is consistent across development. Moreover, developmental changes of occipital alpha are partially mediated by development of the optic radiation. Additional analyses on potential behavioral correlates of alpha confirmed its role for the accuracy of conscious visual target detection^25–28^. This is in line with the idea that the alpha rhythm reflects a general brain mechanism of inhibition that can modulate visual processing by selectively gating the neural signal flow between the thalamus and V1^29–31^. These findings further expand our understanding of alpha by showing a link between the development of visual white matter pathways, brain oscillations and visual behavior.

## Results

### Structural bases of alpha

Individual alpha properties including central frequency, power and bandwidth were estimated with the Fitting Oscillations and One-Over-F (FOOOF) toolbox^32^. These alpha estimates were calculated separately for eyes closed (EC) and eyes open (EO) resting state conditions. DMRI data were processed with QSIprep^33^ and tractometry was performed with pyAFQ^34^ and quality controlled^35^ to identify the optic radiations (and control pathways) in each individual’s brain. We first tested whether alpha features were related to the average fractional anisotropy (FA) of the optic radiations by fitting a linear mixed effects (LME) model on FA that included alpha frequency, power, bandwidth, and age as fixed factors. Site location was included as a random effect (for site effects in the HBN dataset see^35^). Frequency was the only alpha feature that was consistently related to the optic radiations FA across eyes open/closed conditions (frequency, EC: *ß*=0.003, *SE*=0.001, *t*=2.65, *p*=0.008; EO: *ß*=0.002, *SE*=0.001, *t*=2.58, *p*=0.01; power, EC: *ß*=0.008, *SE*=0.003, *t*=2.32, *p*=0.02; EO: *ß*=0.005, *SE*=0.004, *t*=1.06, *p*=0.29; bandwidth: EC: *ß*=0.001, *SE*=0.001, *t*=0.06, *p*=0.95; EO: *ß*=-0.001, *SE*=0.001, *t*=0.71, *p*=0.48). A separate LME model on alpha frequency showed that every increase of 1 Hz in the alpha frequency corresponded to an increase of 0.002 in optic radiations FA after accounting for age and site location (EC: *ß*= 4.26, *SE*= 1.85, *t*= 2.30, *p*=0.021, EO: *ß*= 5.20, *SE*= 2.58, *t*= 2.01, *p*=0.044; Fig. 1). Adding the factor of optic radiations average length did not improve the model fit (starting model, BIC=1836; after adding fiber length BIC=1852), suggesting that individual alpha frequency could be better predicted based on optic radiations FA than fiber length.

**Fig 1.**
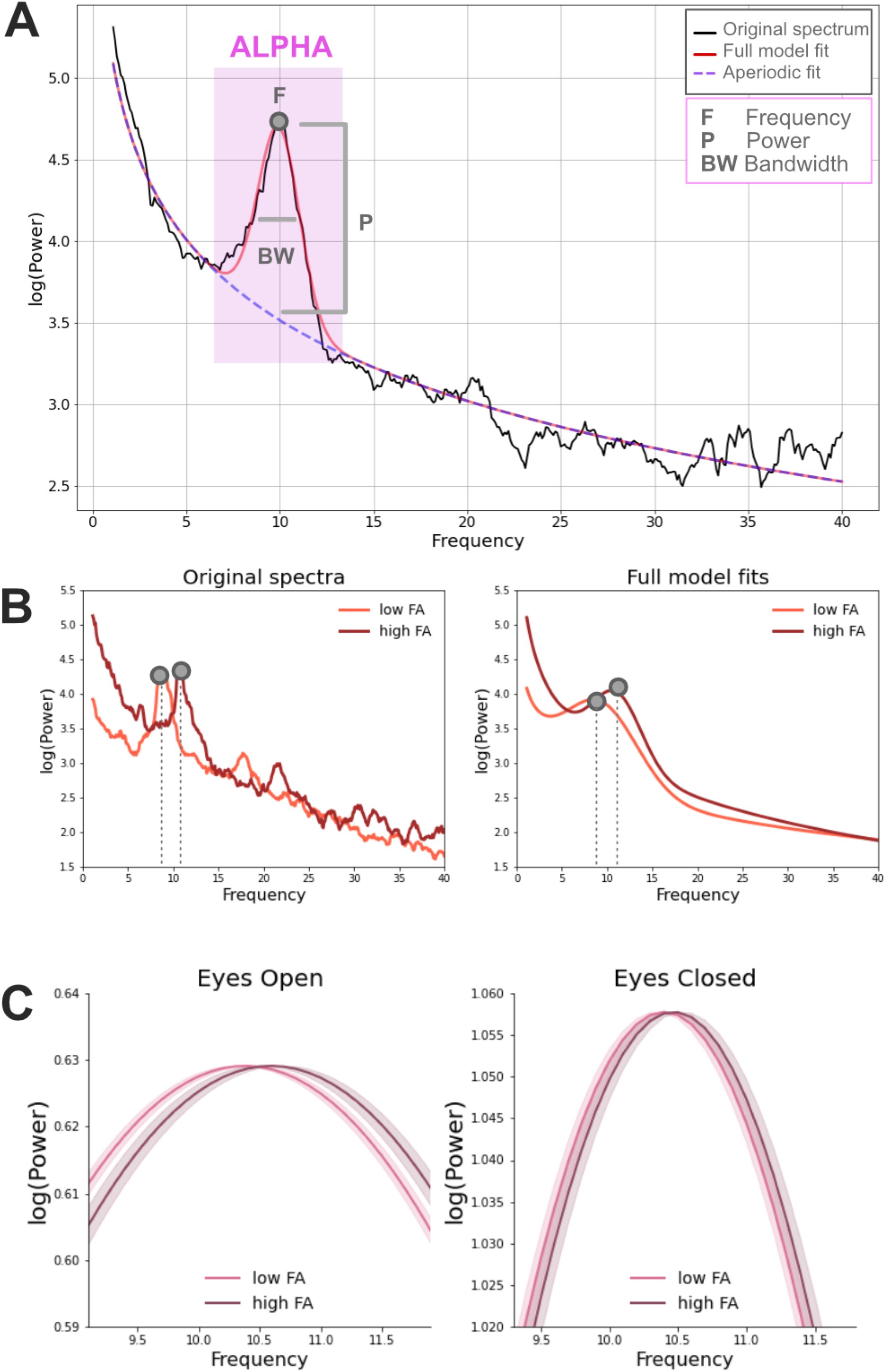
**A:** Example of a power spectrum from a 5-year-old male participant (in black) with closed eyes (EC). The corresponding FOOOF model fit is displayed in red and it corresponds to the sum of the periodic (gaussian function included in the purple square) and the aperiodic signal (dashed line). Three different estimates are extracted from the periodic signal within the alpha frequency range: power, central frequency and bandwidth. **B:** Examples of original power spectra and corresponding FOOOF model fits in the EC condition. Data come from two representative male participants of 12 years of age with high and low FA average values (high FA=0.55: low FA=0.50; median FA=0.53). **C:** Relationship between alpha frequency and the FA of the optic radiations in the EC and EO conditions in the full participant sample. Model fits of the periodic signal are shown for high and low FA participants (defined based on a median split). Beta estimates of alpha frequency were calculated based on the following LME model: alpha ~ FA + age + (1|sj). Model fits of the periodic signal were derived based on the formula 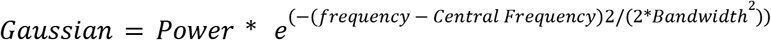. Shaded areas represented +/- 1 SE.

We next examined which part of the optic radiation was related to alpha frequency by fitting the same LME models for each node along the tract profile (n nodes = 100). The LME models included age as a fixed factor and site location as a random factor. The alpha-FA relationship was mainly observed in the centro-posterior part of the optic radiations (Fig. 2; for GAM models results see SI Appendix, Figure S2).

**Fig 2.**
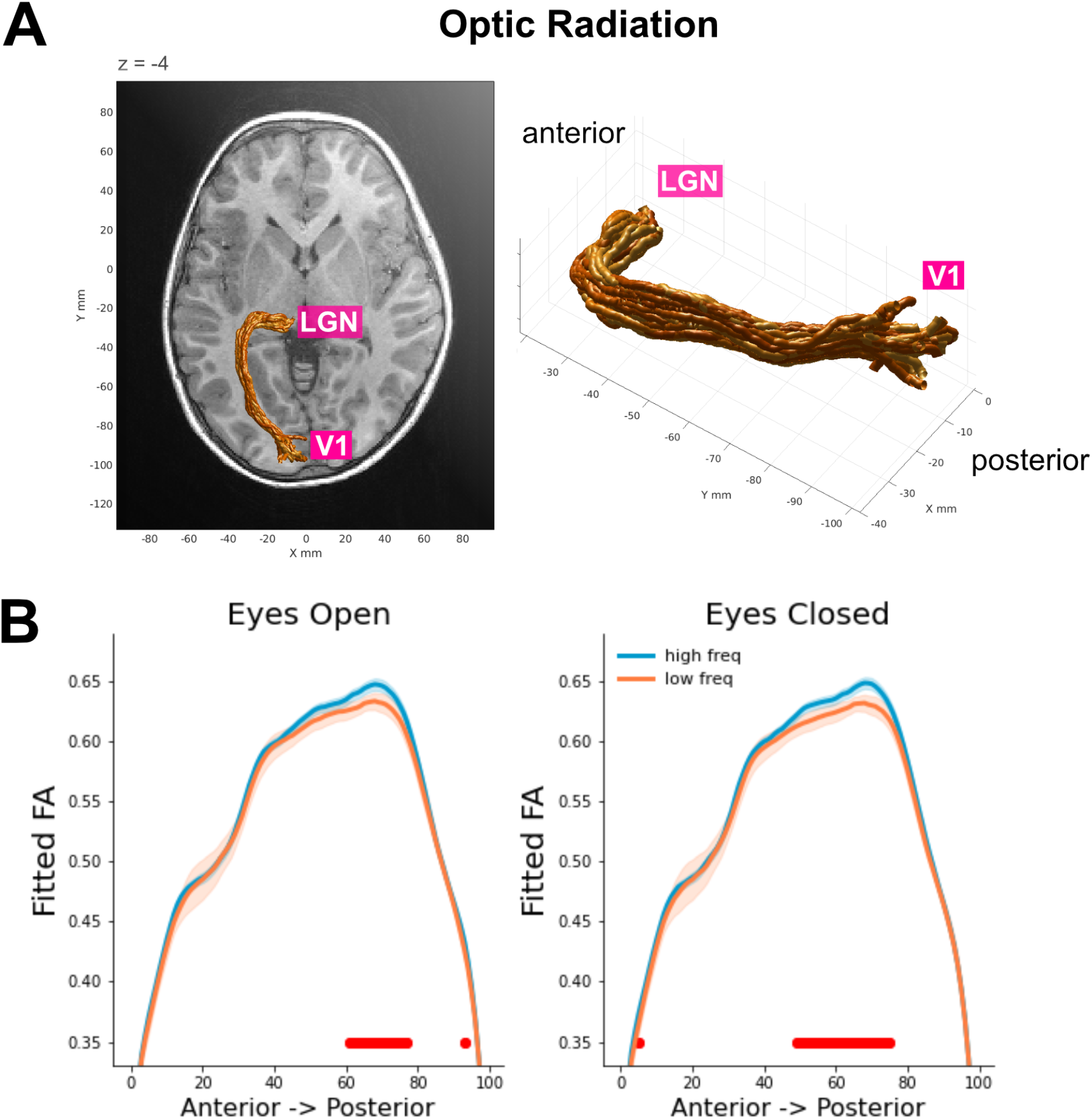
**A:** Three-dimensional rendering of the left optic radiation for a single representative participant (5-year old female). The rendering was derived from the Automated Fiber Quantification software^36^. **B:** Tract profiles of the optic radiations FA for each alpha frequency group (defined based on a median split) in EC and EO conditions. The plots show FA values estimated based on the beta coefficients extracted from node-by-node LME models. The optic radiations FA was modeled as a function of alpha frequency after accounting for age and site location (i.e., FA ~ alpha frequency + age + (1|site)). Red horizontal lines highlight nodes where FDR corrected p-values are below 0.025. Shaded areas represent +/- 1 SE.

We checked for the anatomical specificity of the alpha frequency effect on the optic radiation by running the same LME models on a set of control tracts. We did not see a relationship with alpha frequency in other cortico-thalamic pathways (anterior thalamic radiation: EC: *ß*=1.16, *SE*=2.08, *t*=0.56, *p*=0.58; EO: *ß*=1.58, *SE*=2.76, *t*=0.57, *p*=0.57) or in white matter pathways that end in the posterior part of the cortex (occipital segment of the corpus callosum: EC: *ß*=0.26, *SE*=1.37, *t*=0.19, *p*=0.85; EO: *ß*=3.33, *SE*=2.05, *t*=1.63, *p*=0.10; posterior parietal segment of the corpus callosum: EC: *ß*=0.93, *SE*=1.31, *t*=0.71, *p*=0.48; EO: *ß*=0.32, *SE*=1.61, *t*=0.20, *p*=0.85; inferior fronto-occipital fasciculus: EC: *ß*=0.84, *SE*=1.41, *t*=0.59, *p*=0.55: EO: EO: *ß*=0.53, *SE*=1.72, *t*=0.31, *p*=0.76; Fig. 3).

**Fig 3.**
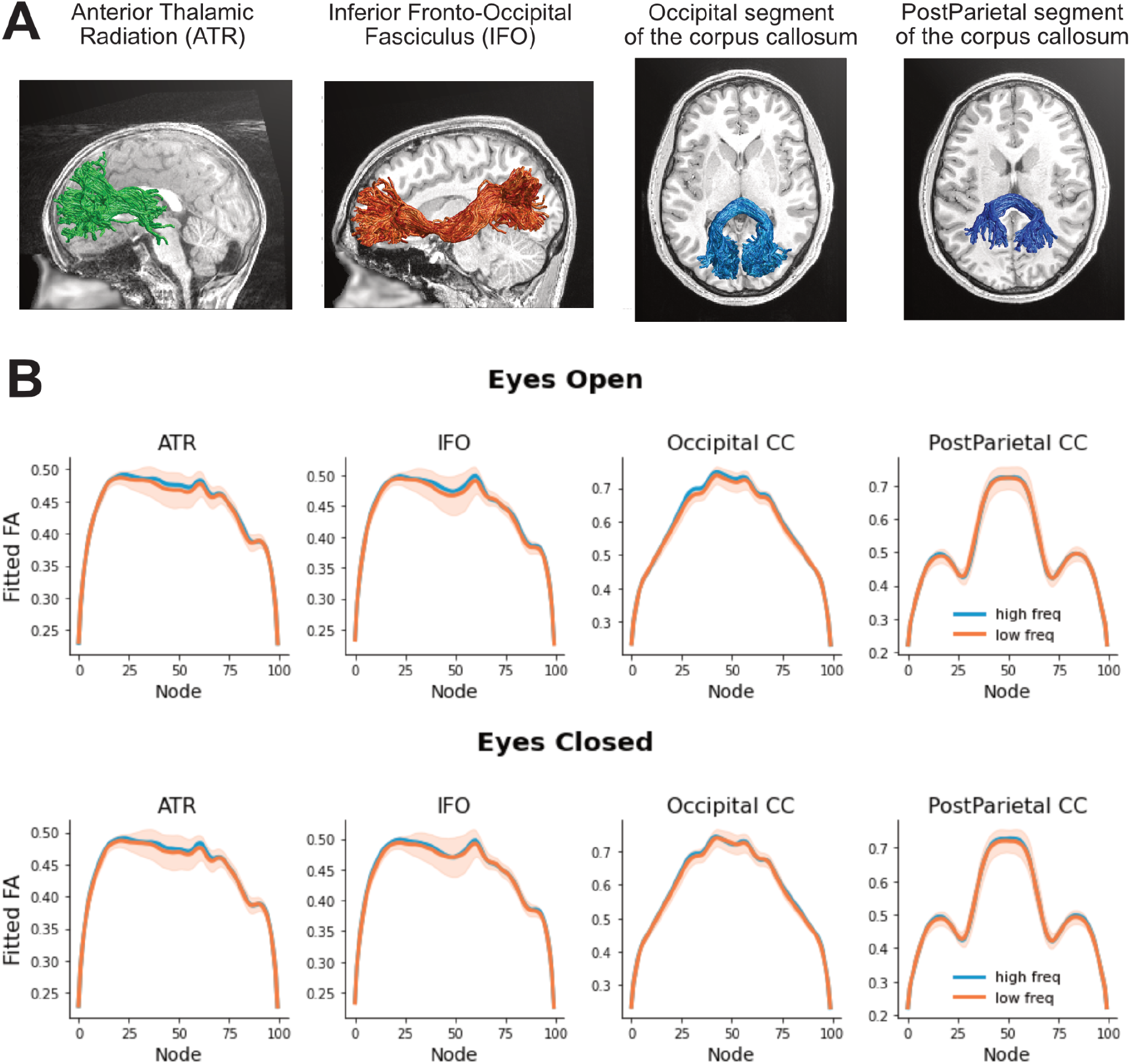
**A:** Three-dimensional rendering of the control tracts from three representative participants (ATR: 6 year-old male; PostParietal: 7 year-old female; IFO and Occipital: 12 year-old male). **B:** Tract profiles of control tracts for each alpha frequency group (defined based on a median split) in EC and EO conditions (first and second row, respectively). The plots show FA values estimated based on the beta coefficients extracted from node-by-node LME models as in Fig. 2. Shaded areas represent +/- 1 SE.

### Other white matter microstructural properties

The FA of the optic radiations was mainly tested as it represents the diffusion property that has been most largely reported to correlate with electrophysiological measures^37–46^. Mean diffusivity measures of the optic radiations did not show a similar relationship with alpha features (frequency, EC: *ß*=-0.002, *SE*=0.001, *t*=1.37, *p*=0.17; EO: *ß*=-0.001, *SE*=0.001, *t*=0.34, *p*=0.74; power, EC: *ß*=0.001, *SE*=0.004, *t*=0.13, *p*=0.90; EO: *ß*=-0.001, *SE*=0.005, *t*=0.10, *p*=0.92; bandwidth: EC: *ß*=-0.001, *SE*=0.001, *t*=1.24, *p*=0.21; EO: *ß*=-0.001, *SE*=0.001, *t*=0.90, *p*=0.37).

### Developmental changes of alpha and optic radiations

A clear developmental trajectory could be observed in alpha estimates and in the optic radiations FA (Fig. 4). Between 5 and 21 years of age, alpha frequency, power and bandwidth increased (frequency, EC: r=+0.36, p=2e-19; EO: r=+0.34, p=1e-17; power, EC: r=+0.18, p=2e-5; EO: r=+0.05, p=0.26; bandwidth, EC: r=+0.17, p=4e-5; EO: r=+0.12, p=0.004; SI Appendix, Figure S3). Moreover, the optic radiations FA increased with age (r=+0.30, p=9e-12).

**Fig 4.**
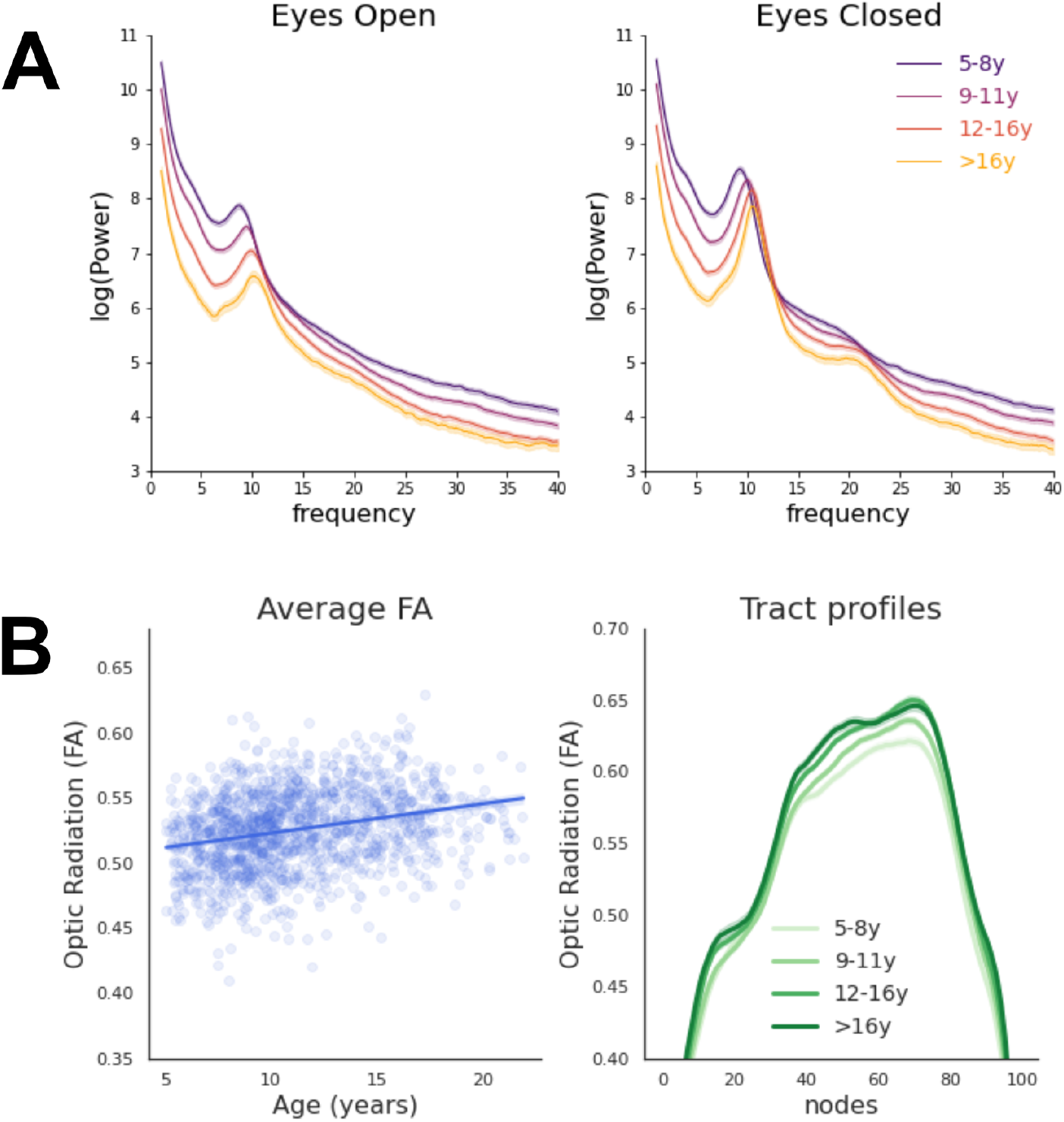
**A:** Developmental trajectory of the power spectra in the EEG final sample (n=1,388) for EC and EO conditions. Age groups approximately correspond to different developmental stages: pre-puberty (5-8 years, n=373), early adolescence (9-11 years, n=488), mid-adolescence (12-16 years, n=416) and late adolescence (>16 years, n=111)^47^. **B:** Developmental trajectory of the optic radiations FA values in the dMRI final sample (n=1,394). Average FA values and FA values along the tract profile are displayed on the left and right panel, respectively. Age groups correspond to pre-puberty (5-8 years, n=308), early adolescence (9-11 years, n=481), mid-adolescence (12-16 years, n=436) and late adolescence (>16 years, n=180)^47^. All shaded areas represent +/- 1 SE.

Despite these robust developmental changes, the alpha-optic radiation relationship was consistent across development. For instance, adding the interaction between age and optic radiations FA did not improve the model fit of the LME model alpha ~ FA + age + (1|siteID) (BIC= 1836; after adding the interaction BIC= 1839), which suggests that the alpha-white matter link was present even after controlling for additive and interactive developmental effects.

We further tested whether developmental changes of the optic radiation mediate development of alpha by performing a causal mediation analysis^48^. Two linear regression models were specified: a mediator model estimating the effect of age on FA, and an outcome model estimating the effect of age and FA on alpha frequency for both EC and EO. The mediation R package^48^ uses these two models as starting points to compute the average causal mediation effect (indirect effect of age on alpha that is related to the FA mediator) and the average direct effect (effect of age on alpha after partialling out the FA mediator effect). The sum of these two effects resulted in the total effect of age on alpha. A bootstrap analysis with 1000 simulations was used to calculate the uncertainty estimates of these effects^49^. This analysis showed that development of the optic radiation partially mediated developmental changes of alpha frequency (EC: average causal mediation effect: *β*=0.008, *CI* [0.009; 0.02], *p*=0.034; percentage of age effect that is due to the FA mediator: 6.44%, *p*=0.034; EO: average causal mediation effect: *β*=0.011, *CI* [0.002; 0.02], *p*=0.02; percentage of age effect that is due to the FA mediator: 6.77%, *p*=0.02; Fig. 5). The effect of age on alpha was still present after taking into account the mediator (EC: average direct effect: *β*=0.11, *CI* [0.09; 0.14], *p*<0.001; EO: average direct effect: *β*=0.15, *CI* [0.12; 0.18], *p*<0.001), suggesting that FA variations partially mediated alpha development.

**Fig. 5.**
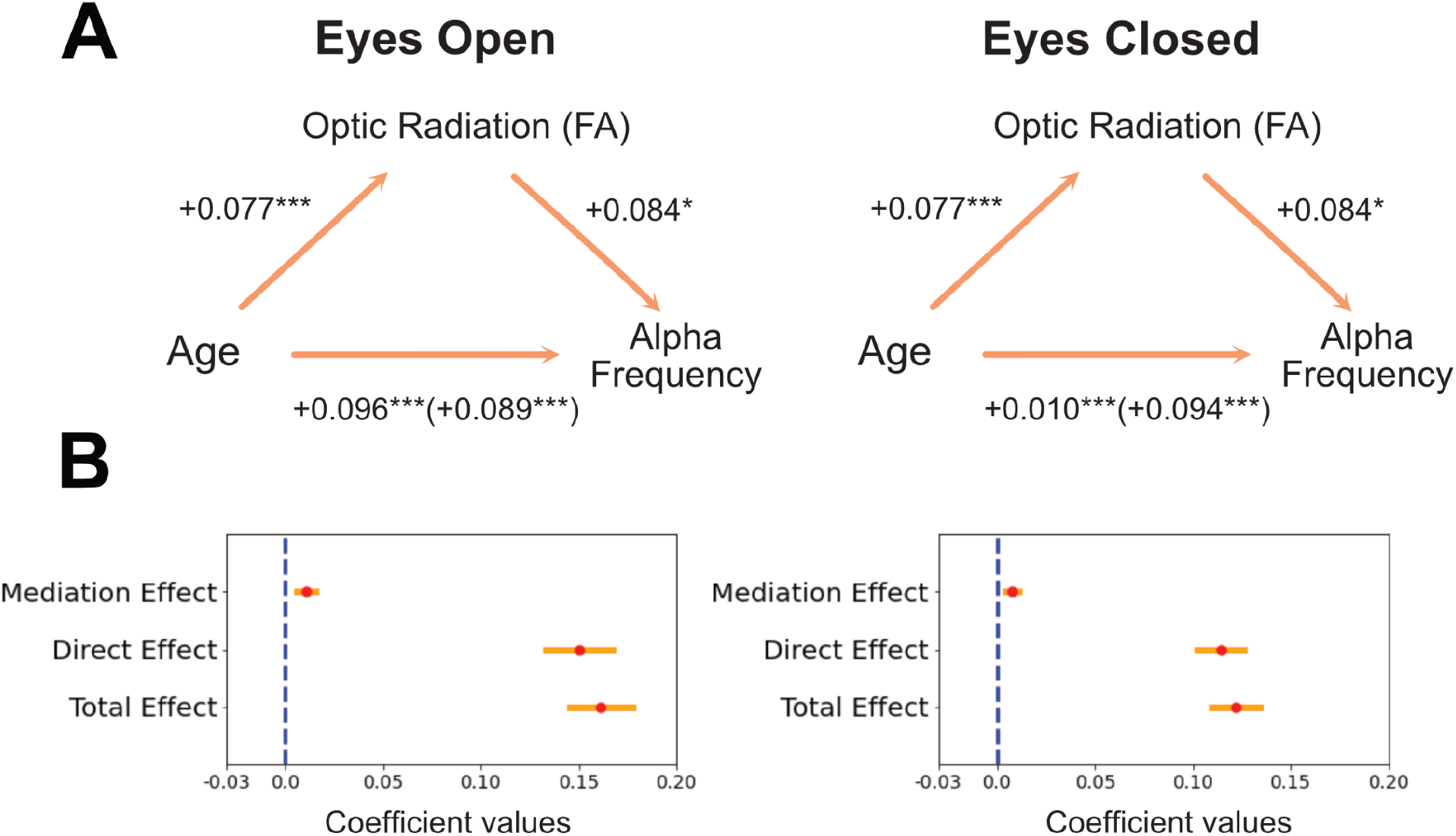
Mediation effect of the optic radiation on alpha development. **A:** Schematic representation of the mediation analysis results. Standardized coefficients are reported. **B:** Beta coefficients of the main effects of the mediation analysis are displayed in red. Yellow bars represent +/- 1SE.

### Behavioral correlates of alpha

Finally, we explored behavioral correlates of occipital alpha by extracting measures of visual perception for each individual. Participants performed a visual contrast change detection task where they were presented with an annular pattern (1° of inner radius; 6° of outer radius), which consisted of two overlaid gratings (each one tilted 45° to the left and to the right, respectively). At the beginning of the trial, these two gratings had the same visual image contrast (50%). Within the following 1600 ms, one of the two gratings gradually changed its contrast to 0% while the other reached a contrast of 100% (total of three blocks with 24 trials each, equally distributed across left and right sides; the experimental paradigm is described in detail in^50^). Participants were asked to press one of two response buttons based on the grating that had the strongest contrast. After 800 ms the gratings’ contrast level came back to baseline (50%) and participants received feedback on their trial performance (ITI could be 2.8, 4.4 or 6 sec). Individual average accuracy scores were calculated and only participants with above-chance performance were considered (n = 917, average accuracy: 83.66, SD: 0.12). A linear regression model was fitted to these accuracy measures including alpha features (power, frequency, bandwidth) and age as factors. Individual alpha frequency was the only electrophysiological property related to contrast detection accuracy (EC: *ß*=0.011, *SE*=0.003, *t*=3.34, *p*<0.001, R^2^=27.7%; EO: *ß*=0.008, *SE*=0.003, *t*=3.18, *p*=0.002, R^2^=27.5%). Drift diffusion models^51^ were also run in order to combine reaction times and accuracy scores within the same dependent variable. Individual drift rates were obtained, which correspond to an estimate of the rate with which the visual system extracts information to inform a decision. The drift rate was associated with alpha frequency (EC: *ß*=0.072, *SE*=0.030, *t*=2.42, *p*=0.016, R^2^=13.8%; EO: *ß*=0.064, *SE*=0.023, *t*=2.70, *p*=0.007, R^2^=13.7%), suggesting that participants with a fast alpha rhythm also extracted visual information more efficiently than participants with a slow alpha rhythm (i.e., they had a high drift rate at the visual detection task). Moreover, a mediation analysis showed that developmental changes of accuracy scores were mediated by alpha frequency changes (EC: average causal mediation effect: *β*=0.002, *CI* [0.001; 0.002], *p*<0.001; percentage of age effect that is due to the alpha mediator: 8.08%, *p*<0.001; EO: average causal mediation effect: *β*=0.001, *CI* [0.001; 0.002], *p*<0.001; percentage of age effect that is due to the alpha mediator: 7.39%, *p*<0.001; Figure 6). Similar mediation effects of alpha were observed with developmental changes of drift rate scores, although only in the EC condition (EC: average causal mediation effect: *β*=0.007, *CI* [0.0003; 0.01], *p*=0.046; percentage of age effect that is due to the alpha mediator: 9.24%, *p*=0.046; EO: average causal mediation effect: *β*=0.006, *CI* [-0.002; 0.01], *p*=0.13).

**Fig. 6.**
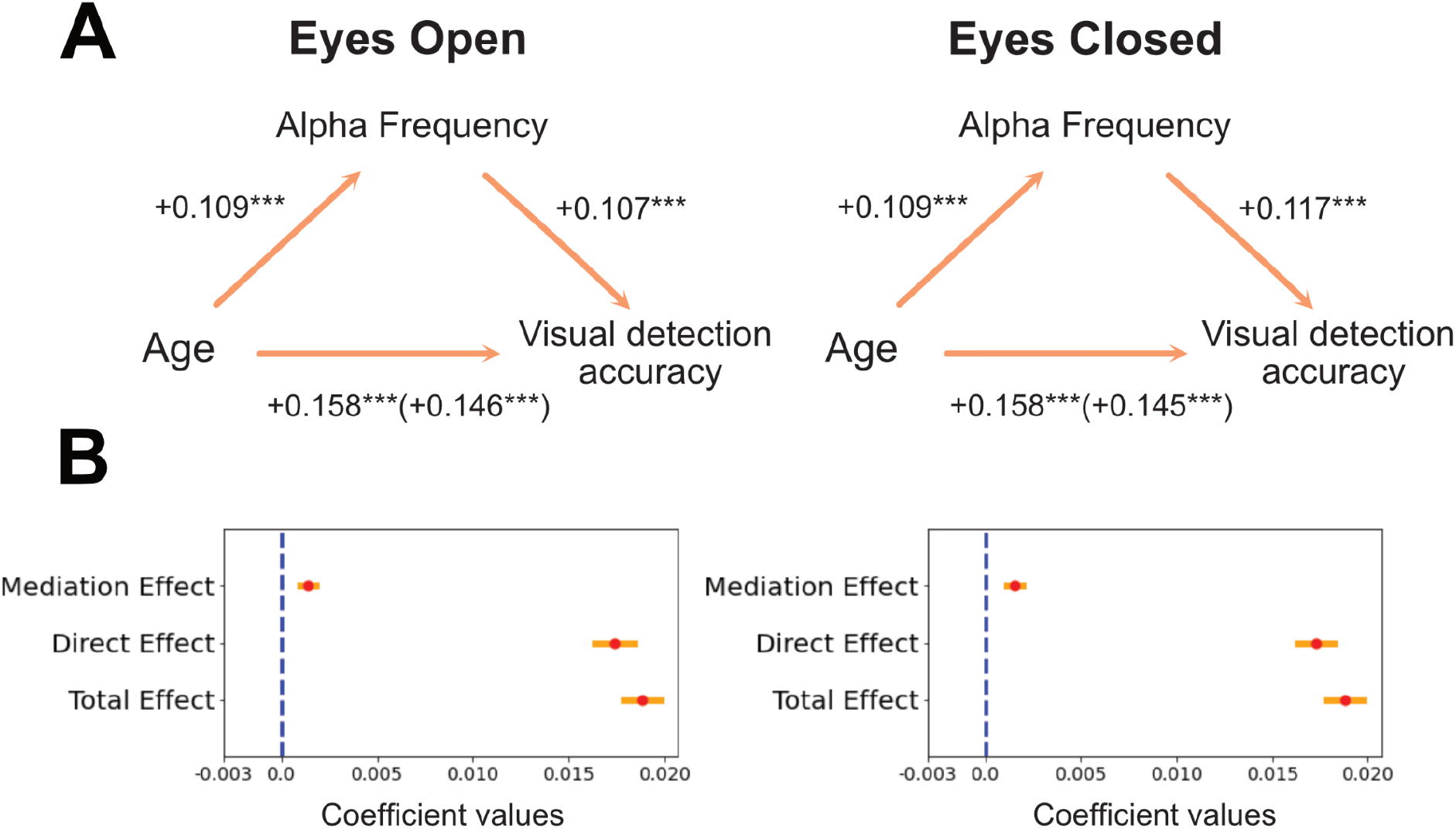
Mediation effect of alpha on visual detection development. **A:** Schematic representation of the mediation analysis results. Standardized coefficients are reported. **B:** Beta coefficients of the main effects of the mediation analysis are displayed in red. Yellow bars represent +/- 1SE.

We finally tested whether the optic radiations FA further contributed to explain the variability in visual perception performances by examining a sample of participants that have EEG, dMRI and behavioral measures available (n = 399). A linear regression model on this smaller dataset confirmed the effect of EC alpha frequency on visual accuracy (EC: *ß*=0.012, *SE*=0.006, *t*=2.04, *p*=0.042, R^2^=26.5%; EO: *ß*=0.005, *SE*=0.004, *t*=1.46, *p*=0.15, R^2^=26.4%), while no effect could be observed for the optic radiations FA (EC: *ß*=0.13, *SE*=0.19, *t*=0.71, *p*=0.48; EO: *ß*=0.13, *SE*=0.19, *t*=0.70, *p*=0.49). No significant effects were observed in the drift rate analysis for this sample.

## Discussion

This work capitalized on a large dMRI-EEG developmental sample to test the relationship between visual white matter pathways and spontaneous alpha rhythm. Our results showed that alpha frequency at rest is specifically related to the structural properties of optic radiations: children and adolescents with a fast alpha rhythm also show high FA values of the optic radiations. This structural-functional relationship was observed after accounting for age effects, suggesting that alpha oscillations have a consistent structural correlate between 5 and 21 years old. Among the optic radiations structural properties that can be at the basis of alpha frequency variations there are axonal density, axonal size, fiber spatial organization, myelination, glial cells structural properties^52,53^(which can all affect FA values^52,54^), while fiber length does not seem to play a crucial role. Interestingly, FA but not MD was related to electrophysiological properties (for a similar asymmetry see^24,40^). This discrepancy might be related to spatial differences in the statistical power of the two methods^55^, as well as to a difference in sensitivity of these two estimates. For instance, fiber orientation coherence is likely to affect FA more than MD^56^ and it also has an impact on neural synchronization, which ultimately modulates the EEG activity measured on the scalp^57,58^. Overall, the observed link between visual white matter microstructural properties and occipital electrophysiological responses supports the idea that the optic radiations FA is related to the coherence with which white matter fibers can deliver neural signals, which are finally reflected by oscillatory brain activity^21,22,44,59^. Future work can identify which microstructural properties mainly contribute to the neural oscillatory modulations observed here.

The portions of white matter fibers that were mainly related to alpha frequency were located in the centro-posterior segment of the optic radiations. There are at least two possible explanations for the location of this effect. First, this segment of the tract represents the closest location to the alpha recording site and cortical generators. Second, the posterior part of the optic radiation has a higher signal-to-noise ratio (and smaller SE) as compared to the anterior segment, which is probably due to the large size of the posterior endpoint ROI (V1) and a more linear trajectory of the posterior as compared to the anterior segment of the tract^20^.

Our findings also showed that development of the optic radiations accounts for changes of alpha frequency between childhood and late adolescence. Between five and 21 years of age the visual brain network undergoes a large range of structural and functional transformations. Visual white matter pathways increase their structural coherence, axonal diameter, axonal density and myelination^60–62^. This happens with a concomitant increase of FA values and improved signal transmission^63^. At the same time, the rhythm of spontaneous occipital alpha speeds up^64^, which probably reflects a higher precision in neural synchrony over long distances and greater coherence between thalamo-cortical alpha generators^65^. Our findings showed that these two maturational phenomena are interrelated, with structural changes of the optic radiations mediating the development of alpha oscillations. This result complements previous reports showing a link between the development of the optic radiations and other types of electrophysiological responses, such as early visual evoked responses peaking around 100 ms^66^. Overall, these findings suggest that the maturation of the optic radiations (and possibly a greater fiber spatial orientation coherence) accounts for changes in the precision and frequency of neural synchronization within the alpha band (i.e., rhythmic neural activity every 100 ms). Future longitudinal studies will help clarify the temporal sequence of these structural and functional changes during development.

Finally, our visual detection task analyses highlighted that a proper development of spontaneous alpha activity has crucial behavioral implications. Data from the contrast detection task confirmed that the individual variability observed in alpha frequency predicts changes in the quality of visual perception. Participants with a fast alpha rhythm showed a more accurate performance at an image contrast detection task and a more efficient visual information extraction. These results are fully in line with previous reports showing that alpha speed determines the temporal resolution at which visual information can be consciously sampled^27^, as well as studies showing a link between prestimulus alpha and conscious visual detection^26,28,67,68^. All these findings are compatible with the hypothesis that alpha oscillations represent a general electrophysiological mechanism of rhythmic inhibition pulses (every 100 ms) that is able to cyclically modulate the level of excitability of a given brain area^29,69^ (e.g. visual cortex). According to this perspective, spontaneous occipital alpha reflects a general gating mechanism that regulates neural information flow between the thalamus and the visual cortex and ultimately impacts our conscious visual perception. Our findings showed that the maturation of the optic radiation mediates the development of this inhibition mechanism reflected by alpha, which ultimately contributes to visual detection accuracy improvement during childhood and adolescence.

In summary, alpha is a predominant rhythm of our brain and its characteristics can widely vary over development. Individual variability in alpha frequency is specifically related to the structural properties of visual white matter pathways and can ultimately predict the rate of our visual information extraction. This work shows that the maturation of optic radiations is linked to an increase of alpha frequency, which contributes to visual detection enhancement over childhood and adolescence.

## Methods Participants

Participants data came from the Healthy Brain Network pediatric mental health study (HBN^70^), which include dMRI (initial raw data sample size = 2,747) and EEG (initial raw data sample size = 2,561) measures of English speaking children and adolescents from 5 to 21 years of age. Exclusion criteria included: moderate to severe cognitive impairment (IQ below 65), encephalopathy, neurodegenerative disorders, hearing and visual impairment All participants had normal or corrected to normal vision. Informed consent was obtained for each participant above 18 years of age. Written assent was obtained for younger participants, and their legal guardians were asked to sign a written consent.

### dMRI acquisition and preprocessing

Diffusion MRI (dMRI) data were acquired using a 1.5T Siemens mobile scanner and three fixed 3T Siemens scanners in the New York area (four locations: Staten Island, Rutgers University Brain Imaging Center, the CitiGroup Cornell Brain Imaging Center, and the City University of New York Advanced Science Research Center). Voxel resolution was 1.8×1.8×1.8 mm with 64 non-collinear directions measured for each of b = 1000 s/mm2 and b = 2000 s/mm2. QSIPrep^33^ preprocessed dMRI data were accessed from AWS S3 at s3://fcp-indi/data/Projects/HBN/BIDS_curated/derivatives/qsiprep/ together with individual quality control scores (n=1885, for the preprocessing pipeline description and quality control scores definition see^35^). The left and the right optic radiations were identified using pyAFQ^34^ based on two endpoint regions of interest (ROIs, for a similar pipeline see^66^): the primary visual cortex and the central part of the thalamus including the lateral geniculate nucleus (defined based on the AICHA atlas,^71^; minimum distance 3 mm). Three exclusion ROIs were also used to further clean the tract from crossing fibers (temporal pole, and occipital pole from the AICHA atlas, and the posterior portion of the thalamus based on the brainnetome atlas; minimum distance 3 mm^20,72^). All ROIs were defined in a MNI template and transformed to each participant’s native space. A final cleaning step was carried out to remove outlier fibers based on streamline average length and mean Gaussian distance from the bundle core (distance threshold: 3 mm; length threshold: 4 SD^36^). Diffusion metrics were projected onto the optic radiations and fractional anisotropy (FA) was mapped onto each tract, weighting the values based on the streamline’s distance from the core of the tract^36^. Overall, we could detect 1,798 left optic radiations and 1,799 right optic radiations. Further analysis was conducted only on participants where both left and right optic radiation could be found (n=1,774). FA values of the left and right optic radiations were averaged for each participant. Moreover, the mean length of the optic radiations was calculated by averaging the median values of the streamlines length for the left and right optic radiations (step size=0.5 mm). A similar pyAFQ pipeline was used to segment default white matter bundles (total: 24), among which a subset of cortico-thalamic (anterior thalamic radiation and corticospinal tract) and posterior bundles (inferior fronto-occipital fascicle, occipital and posterior parietal part of the corpus callosum) were used as control tracts. Only participants with quality control scores higher than 0.3 (based on^35^) were included in the final statistical analyses (final dMRI sample size=1,394).

### EEG acquisition and preprocessing

EEG data were recorded with a 128-channel EEG geodesic hydrocel (Magstim EGI) in a sound-shielded room of one of the four New York recording locations (sampling rate=500 Hz; bandpass filter=0.1-100 Hz). The online reference was at Cz. The impedance was kept below 40 kOhm and tested every 30 min of EEG recording. EEG raw data were accessed from s3://fcp-indi/data/Projects/HBN/EEG.

EEG analysis was performed on a cluster of 13 occipital electrodes (E69, E70, E71, E72, E73, E74, E75, E76, E81, E82, E83, E88, E89) using MNE^73^. The EEG signal was re-referenced offline to the mastoids’ average activity. High and low pass filters were applied (1 and 40 Hz, respectively). Epochs of 10 sec were segmented for each condition (tot 10 epochs, 5 for EC and 5 for EO condition). Bad EEG epochs were automatically rejected or corrected by using the autoreject algorithm^74^, which has been already employed for the preprocessing of big electrophysiological dataset (Human Connectome Project^74,75^). Having an additional preprocessing step where ocular artifacts were further corrected through ICA did not affect the structural-functional results reported here (SI Appendix, Figure S3). Only participants with at least two clean epochs for EC and EO (minimum duration of clean EEG signal = 20 sec) were further analyzed (n=2,364). A trial-by-trial time frequency analysis was performed using a multitaper estimation of the power spectra^76^, then averaged across epochs and electrodes of the cluster. The power spectrum was estimated from 1 to 40 Hz, with a window half-bandwidth of 4 Hz. We used the Fitting Oscillations and One-Over-F (FOOOF) toolbox^32^ to estimate the periodic and aperiodic signals of each individual power spectrum (peak width=0.5-20; maximum oscillatory peaks=1; minimum peak height=0.2). Most of the participants had a maximum power peak around 10 Hz in both EC and EO conditions (SI Appendix, Figure S5). Only those participants that showed a maximum oscillatory peak in both EC and EO within 5 and 15 Hz were included in the next steps of the analysis (n=1,820). The final EEG sample included only subjects where FOOOF models accurately fit the individual power spectrum (r^2^>0.75; final EEG sample size=1,388). Alpha power, frequency and bandwidth were estimated for each participant and condition. The relationship between alpha features and white matter was tested after intersecting the dMRI and EEG samples (final EEG-dMRI sample size n=585).

## SI Appendix

**Figure S1.**
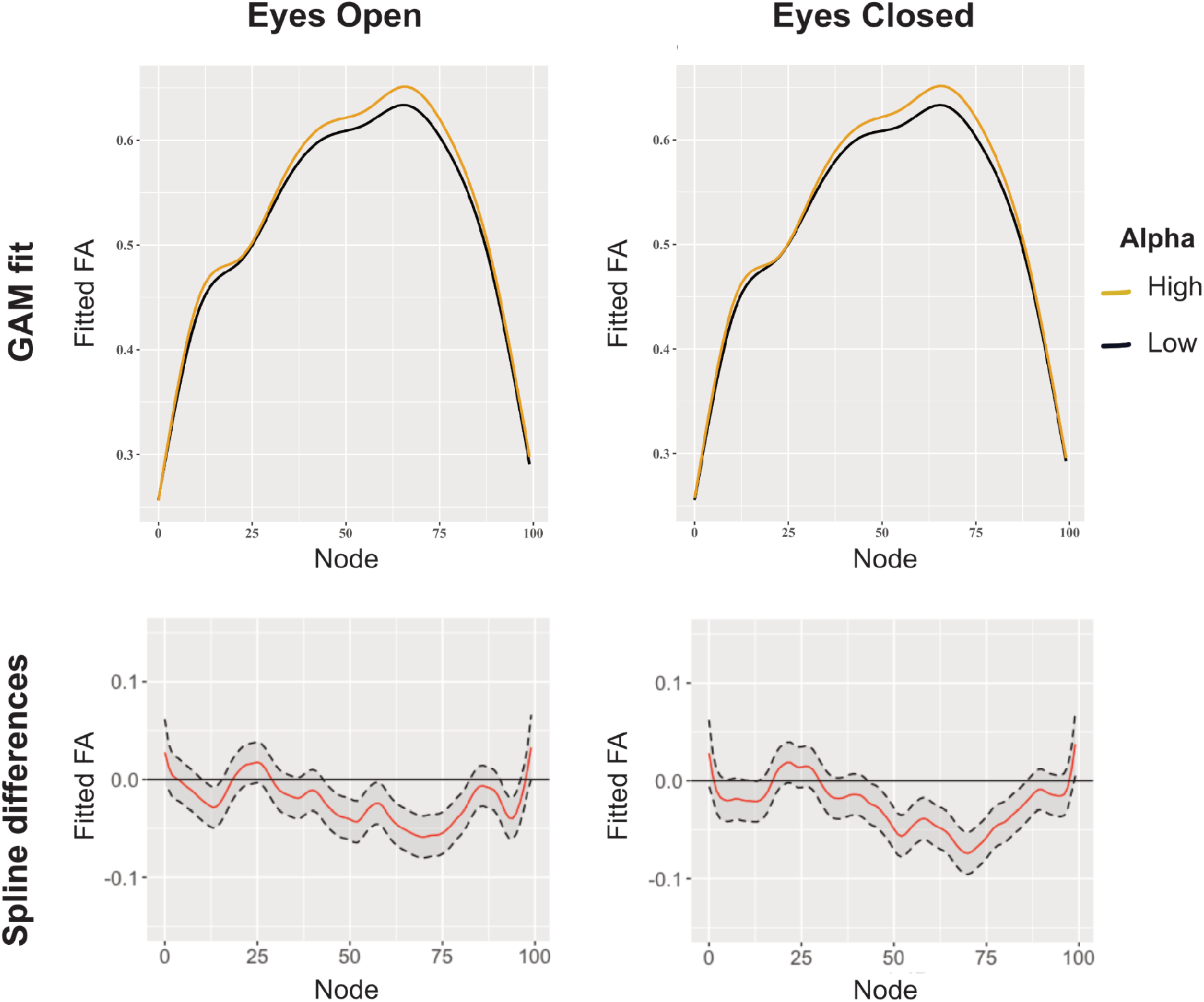
GAM results for tract profile analyses. We adopted a multi-analysis approach where both LME and GAM models were applied for tract profile analyses (following these guidelines ^77^). GAM models were run on the dMRI-EEG sample (n=585) using the tractr R package (https://github.com/richford/tractr), which implements the pipeline reported in ^78^. Our GAM formula was: FA ~ age + alpha frequency + s(nodeID, by = alpha frequency, k = 64) + s(subjectID, bs = “re”). Similarly to what seen in the results from the node-by-node LME models, GAM analyses showed that alpha frequency had an impact on the optic radiations FA and this effect was mainly localized in the centro-posterior part of the tract (EO: estimate= 0.021, SE= 0.002, t= 12.03; EC: estimate= 0.024, SE= 0.002, t= 13.73).

**Figure S2.**
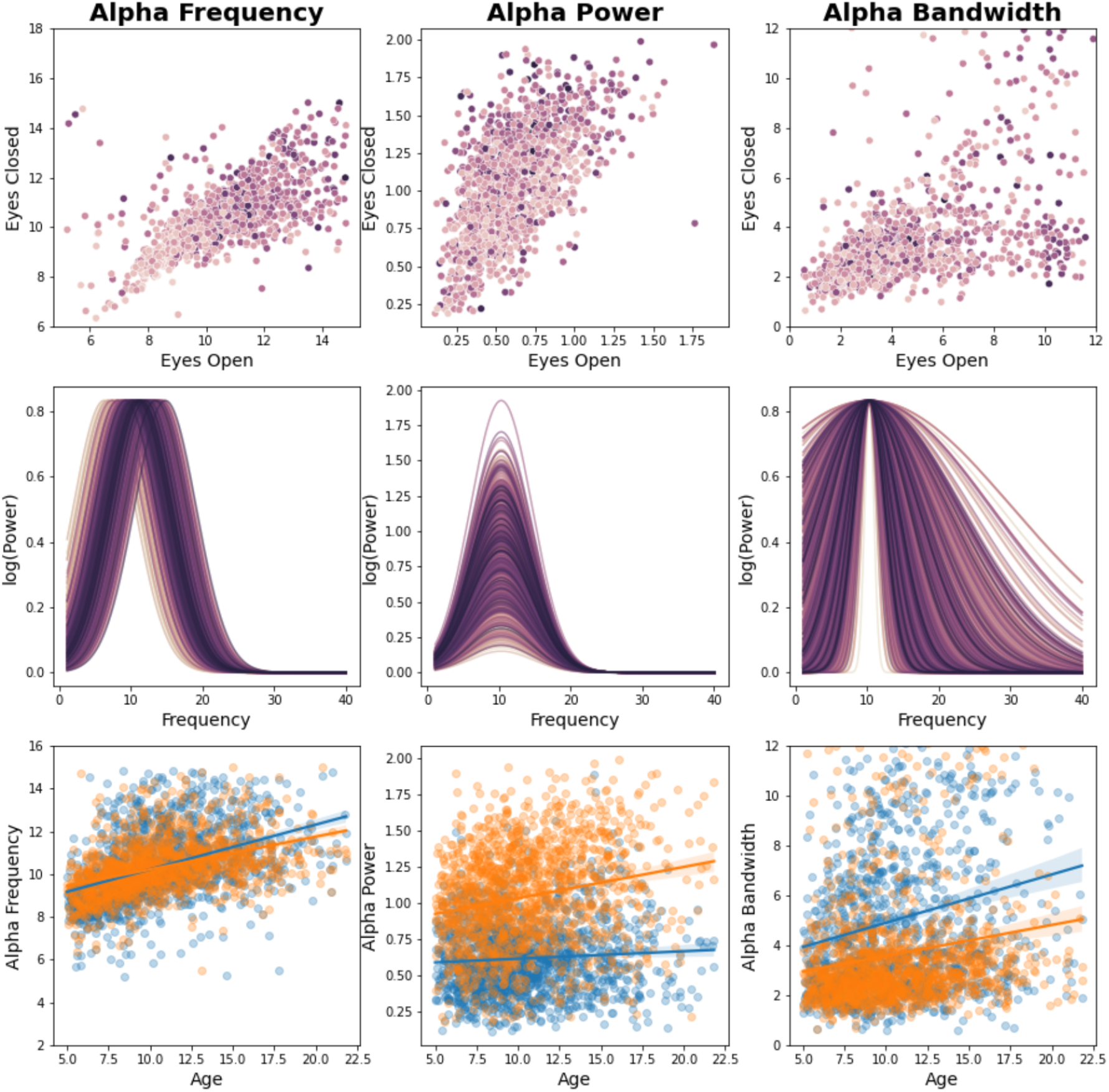
Developmental effects on EC and EO alpha Individual alpha properties (frequency, power and bandwidth) are shown for each condition (EC and EO). The first row shows that alpha frequency and power in EC highly correlate with alpha frequency and power in EO, suggesting that a similar electrophysiological phenomenon is observed across conditions. The second row shows the effect of age on individual FOOOF^32^ models of alpha (calculated as 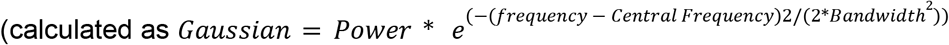). The third row shows the relationships between alpha features and age for both EC and EO conditions.

**Figure S3.**
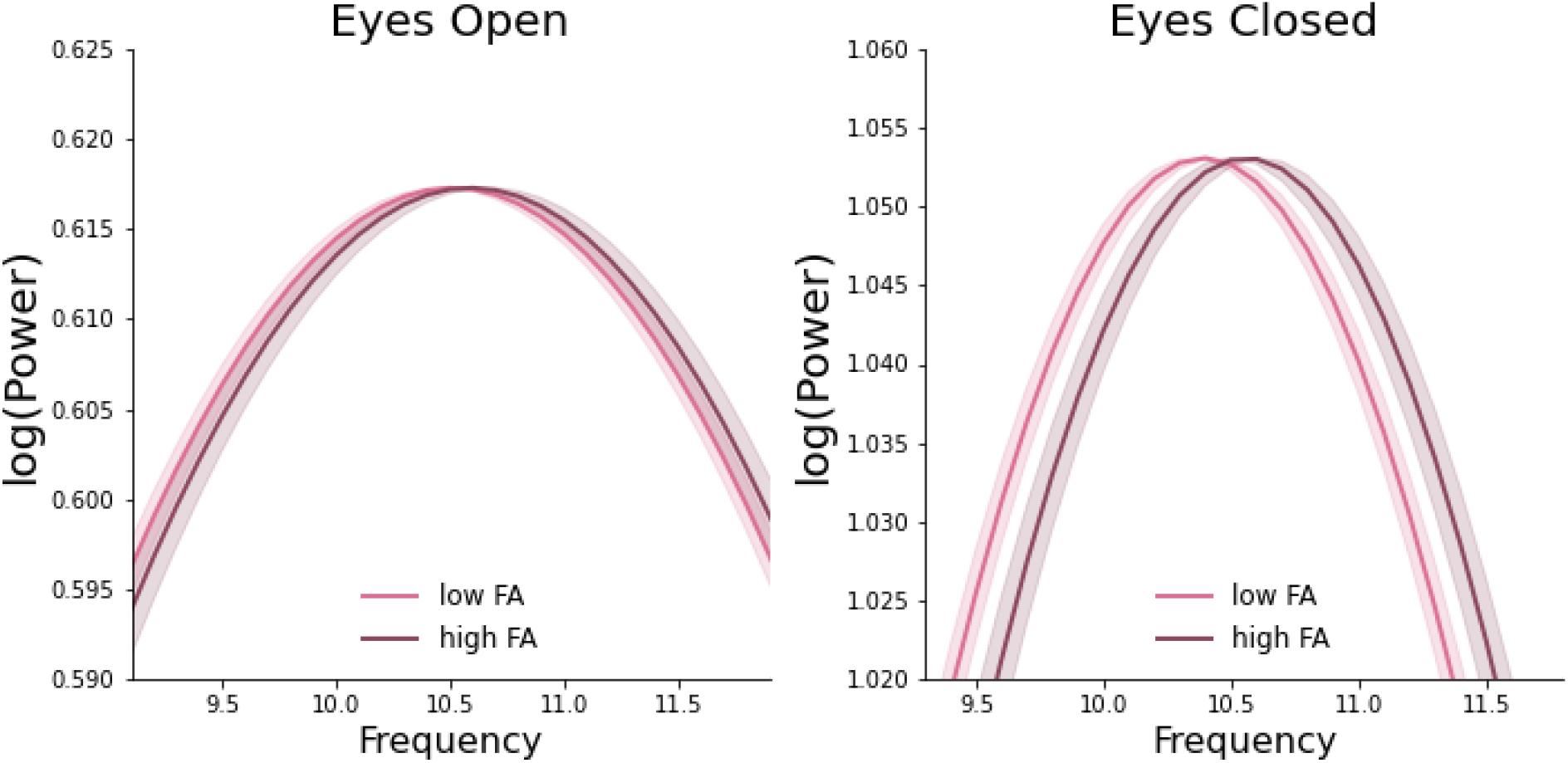
After adding ICA to the EEG preprocessing pipeline the alpha-white matter relationship is still observed. A different EEG preprocessing pipeline was also performed to make sure that ocular movements did not affect the present findings. An Independent Component Analysis (ICA) was applied to the filtered EEG signal obtained as described in the section “**EEG acquisition and preprocessing**”. Ocular artifact detection was based on Pearson correlations between the ICA components and the filtered electro-oculogram (EOG) channels (n=6). Thresholding was based on adaptive z-scoring where z-scores were recomputed until there was no component exceeding the threshold (z-score threshold=3). Independent components that were highly correlated to electro-oculogram channels were excluded. The cleaned EEG signal was epoched and further analyzed as described in the main text. LME models on these EEG data showed again a relationship between alpha and visual white matter pathways. Individual frequency was related to the optic radiations FA. This effect was more evident in the EC condition, which has the highest signal to noise ratio (frequency, EC: *ß*=0.003, *SE*=0.001, *t*=3.33, *p*=0.001; EO: *ß*=0.001, *SE*=0.001, *t*=1.87, *p*=0.06; power, EC: *ß*=0.009, *SE*=0.003, *t*=2.67, *p*=0.008; EO: *ß*=0.006, *SE*=0.004, *t*=1.39, *p*=0.17; bandwidth: EC: *ß*=0.001, *SE*=0.001, *t*=0.46, *p*=0.65; EO: *ß*=0.001, *SE*=0.001, *t*=0.32, *p*=0.75). A separate LME models on alpha frequency showed that a higher alpha rhythm corresponded to an increase of optic radiations FA after accounting for age and site location (EC: *ß*= 5.32, *SE*= 1.80, *t*= 2.96, *p*=0.003, EO: *ß*= 4.68, *SE*= 2.26, *t*= 2.07, *p*=0.04; Fig. S3).

**Figure S4.**
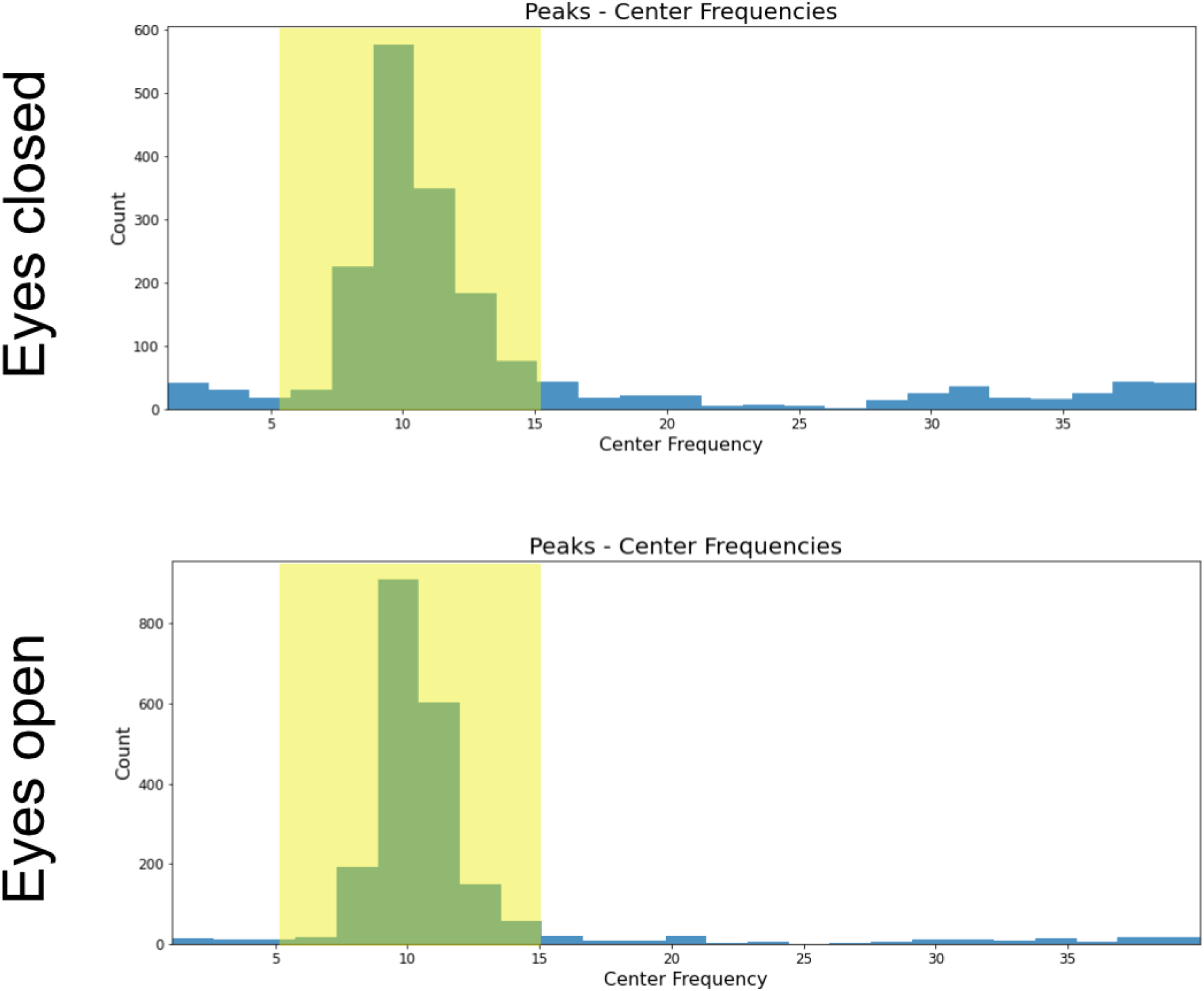
Most participants had a maximum peak within alpha frequency range for both EO and EC conditions.

